# Adaptive Artificial Intelligence to Teach Interactive Molecular Dynamics in the Context of Human-Computer Interaction

**DOI:** 10.1101/2023.08.26.554965

**Authors:** Mustafa Demir, Sean M. Leahy, Punya Mishra, Chun Kit Chen, Abhishek Singharoy

## Abstract

Artificial Intelligence (AI) can be easily integrated into virtual education to drive adaptive instruction and real-time constructive feedback to students, offering a possible conduit for fostering discovery curiosity in learners. This study examines and characterizes Human-AI-Teaming (HAT) coordination dynamics to monitor the inception of discovery curiosity in online laboratories of interactive molecular dynamics (IMD). We used molecular physics measures (kinetic/ potential energy and action) obtained from simple and complex examples of simulated mouse tracking datasets in IMD log files as a proxy for understanding the context of molecular sciences and developing novel interactions for inquiry. These measures are good features of our HAT context because kinetic energy reflects the system’s atoms’ overall motion regarding the individual atoms’ speed. While kinetic energy represents if a learner applies artificial forces to the task, potential energy can be AI’s response to these forces. The action is a systems-level reaction to the changes during the task. By applying nonlinear dynamical systems methods to the physics measures, we extracted the Largest Lyapunov Exponent and Determinism metrics as HATs’ coordination stability and predictability, respectively. The findings underline that while the more complex IMD task required less stable and predictable HAT coordination dynamics, the simple task is more. One explanation is that AI needs to anticipate the learner by providing feedback at the right time and place during the more complex IMD task to initiate and sustain the learner’s discovery curiosity. In IMD, future HAT design should consider coordination dynamics for fostering ‘discovery curiosity’ and practical learning.

## I. Introduction

Artificial Intelligence (AI)’s omnipresence is already rich in various industries worldwide. In the education industry, AI can be integrated into virtual education to drive adaptive instruction and real-time constructive feedback from students, offering a possible conduit for fostering ‘epistemic curiosity’ in learners [1]. Epistemic curiosity is an emotional-motivational state to obtain new knowledge “that motivates individuals to learn new ideas, eliminate information - gaps, and solve intellectual problems” ([2]; see [4]). Even the most advanced students often struggle to understand complex foundational ideas and concepts in a discipline, which is a barrier to asking productive questions, getting in the way of deeper learning. There are two divergent forms of epistemic curiosity, namely the aforementioned ‘discovery curiosity,’ which generates a positive effect when learning something new towards exploratory or mastery-oriented learning [1]–[3], and ‘deprivation curiosity,’ which “involves the reduction of uncertainty, specific exploration, acquiring information that is missing from an existing knowledge-set and performance-oriented learning” [2]. However, ‘deprivation curiosity’ is also associated with errors, confusion, and lack of humility as well as falling for fake news [1]. Thus, this study mainly discusses discovery curiosity for preparing the next generation of AI-powered K-12 technologies in the context of human-computer interaction (HCI).

To do so, we track to monitor the inception of curiosity in online laboratories of interactive molecular visualization [5]. Such virtual labs are prepared to overcome the presentational limitations of textbooks to train students in foundational biotech concepts, including structure and dynamics, sensing, genetics, photosynthesis, and vaccines, with content aligned to standards. Data from the interactive molecular dynamics (IMD) simulations performed in labs offer “real-time” monitoring of protein movements using so-called molecular dynamics (MD) simulation. This data accrued over the interactions offers an opportunity to evaluate whether specific types of HCI during IMD simulations can lead to the elements of discovery curiosity. Specific signatures of HCI are needed to classify the merits (or demerits) of a particular interactive move during molecular dynamics, which we now analyze in real-time metrics that are extracted from nonlinear dynamical systems methods (NDS).

In this study, we applied NDS methods on molecular physics measures (classical energy and action [6]) from mouse tracking datasets in IMD log files as a proxy for understanding the context of molecular sciences and developing novel interactions for inquiry. From these methods, we extracted two NDS metrics: (1) the ‘Largest Lyapunov Exponent’ or stability and (2) ‘determinism’ or predictability [7]. We discuss the findings from these metrics and their visual constructions by considering changes in IMD trajectories as a signature for HCI. We believe these initial findings can guide the design of AI-tutoring technologies in molecular modeling tasks of future virtual classrooms, laboratories, and other high-tech learning environments.

### A. Human-Computer Interaction and Human-AI Teaming

With natural language processing, computer vision, and Machine Learning (ML) advancements, AI has empowered valuable, intuitive interfaces with human-centered design. While AI can offer personalization with reinforcement and evaluation of the interaction between system components, HCI comprises a much bigger picture of an interactive system by emphasizing the user needs, AI design, and development with an optimum level of transparency and explanations, linking human user and AI collaboratively to accomplish a relevant task [8]. Therefore, AI nested (within HCI) greatly benefits human users as long as they are designed to adapt to the user and effectively work in a taskwork environment.

Within the HCI concept, if at least one human and AI agent communicate and coordinate to accomplish a common goal or task, this socio-technical system is called a human-AI teaming (HAT) [9]. In order to consider an AI agent as a true team member within a HAT, it has a task-performing role that is heterogeneous from, but interdependent with, those of other team members, in line with a common goal or task [10], and entails two further requirements. First, if AI serves as a “team member,” it may also need to interact (communicate and coordinate) with other teammates (humans or AI) effectively in a limited time based on their taskwork and teamwork requirements. Second, AI must also be able to work alongside human or nonhuman counterparts and carry out the fundamentals of taskwork and teamwork. An AI teammate needs to demonstrate behavior that must thus be able to respond to situations that may not be anticipated or programmable in their designs. The behavior adapts to changes in a complex task environment in which the system (team) is subject [11].

### B. Discovery Curiosity in HAT

A dictionary definition of curiosity is “an eager wish to know or learn about something” that is rare and unusual [12]. In psychology, curiosity is a stimulus to minimize or eliminate an uncertain state or process by which exposure to novel and complex stimuli occurs. Minimizing or eliminating uncertainty could be achieved by obtaining new knowledge about such stimuli and rewarding the individual [13].

In a traditional learning environment, a learner seeks an instructor’s explanation about a complex learning task [14] that is driven by epistemic curiosity, i.e., a motivated desire to obtain new knowledge during the learning process [1], [2], [13], [15]. Thus, a learner’s epistemic curiosity can be activated when there are expectations of discovery opportunities for learning new knowledge (via the discovery or gathering of new information) that would generate pleasurable experiences of situational interest [7]. This general tendency to delight in gaining new information leads to “greater general knowledge, enhanced responsiveness to new information, and superior ability to distinguish between real and made-up concepts, all complimented by a penchant for intellectual humility” [1]–[3]. Discovery curiosity is a cognitive construct that encourages people to acquire new knowledge and understanding by encompassing a host of virtues: “stimulating positive affect, diverse exploration, learning something completely new, and mastery-oriented learning” [1]. However, there are also possibilities that curiosity can lead to incorrect information and, in turn, obtaining wrong knowledge (see Darker-Side of Curiosity [3]). For instance, during the learning process, a learner can have “feelings of uncertainty and tension that motivate information-seeking and problem-solving behavior,” deprivation-curiosity [6] can result in fake news and wrong information [1], [2], [13] because of negativity of the situation. This study specifically focuses on ‘discovery curiosity’ to avoid its darker side.

In a team context, epistemic curiosity and learning can be indicated by who knows what within a team [16]; they can be achieved via interaction, i.e., coordination of information exchange among the agents (either human or AI). To this end, in engineering, it is assumed that AI also has some level of curiosity, which alters maximizing the AI’s learning about itself and its team task environment by selecting optimum-level actions. In this manner, AI’s artificial curiosity is ‘driven’ by intrinsic motivation for maximum learning that engages reinforcement learning, in which intrinsic rewards are proportional to their learning progress [17], [18].

Compared to an individual level, the HAT information-seeking and problem-solving [19] behaviors that emerged from interactions among the team members might be more effective in obtaining new knowledge to minimize or eliminate the uncertainty. In this case, ‘discovery curiosity’ at an individual and team level during problem-solving and their learning outcomes are influenced by team coordination dynamics to obtain new taskwork or teamwork knowledge [20]. Effective team learning requires establishing a dialogical space between the team members to balance team coordination, i.e., stable and predictable coordination dynamics at some level, leading team members to understand the problem, evaluate possible solutions, and construct knowledge [12,13]. To better understand this balance in HAT, this study uses interactive molecular dynamics within the context of HCI.

### C. Interactive Molecular Dynamics (IMD)

#### 1) Definition

IMD is a computational simulation technique [22] grounded on molecular dynamics simulations, which simulate molecular systems using classical and quantum mechanics or a combination of both, often with parameters extracted from experimental data or quantum chemistry calculations [23]. IMD enables researchers to study the dynamic behavior of molecular systems in real-time by manipulating and visualizing the underlying simulations interactively using a computer interface [22], [24], [25] with sufficient computing power.

IMD’s ability to interact with its users makes it an advantageous technique for educational purposes, as it provides instant feedback to student(s)’ questions, which are being asked through manipulating a molecular dynamics simulation. Given that IMD uses real-world scientific data, the feedback that IMD provides for students is the best “approximate” guess a scientist can provide from the currently available research results and underlying model for the simulations; thus, the educational materials and the available computing resources for the students. Therefore, the IMD’s feedback that a student receives is scientifically relevant yet can be constantly refined by employing additional research data, adopting newer state-of-the-art models, and accessing more computational resources. As a result, a student does not just obtain a simple, schematic understanding of a scientific subject or a molecular system, but rather a first-hand experience with actual scientific research. Thus, if students are willing to and if resources allow, students can constantly ask more questions to improve their models and have updated responses nearly immediately. It is assumed that this research-question-response cycle can likely drive students’ interest (and discovery curiosity) in a scientific question and lead to spontaneous learning of a particular subject. Such spontaneity and increased curiosity is the goal of this project and proof-of-concept study.

#### 2) Simulated IMD Task

To illustrate and characterize this process, we focus on protein folding, a complex process that occurs at a timescale of microseconds to milliseconds [26]– [30]. This timescale is generally prohibitive for molecular dynamics simulations due to its exceptionally high demand for computing power [29], [30]. We have, therefore, decided to have students learn about protein folding by unfolding a native protein instead of folding a purely 1-dimensional amino sequence from scratch. This approach has several advantages. First, unfolding a protein is computationally cheap and quick as a learning task. Students can apply multiple sets of forces to speed up an unfolding process, and any partially unfolded state (several despite being hypothetical) achieved is a successful attempt. Though the unfolding pathways using IMD are not expected to be correct, such partial unfolding of a protein can inform students about concepts of the underlying ‘free energy funnel’ [26] landscape of the protein along the set of actions that they perform, which explains how likely a protein can naturally fold along the steps they attempt. Secondly, students can make multiple attempts with various unfolding strategies within the computational resources allowed. These attempts can be made on the native proteins or on some partially unfolded states. With these flexibilities, the increased number of attempts will facilitate students’ freedom to explore the phrase space of protein folding, subsequently increasing the chance to make discoveries. Lastly, students are exposed to more rare high-energy protein structures called ‘intermediates.’ This increased exposure is intended to raise the likelihood that students will question the observations they make and subsequently stimulate their curiosity of protein structures and movements, their biding to ligands and love vs hate for water (hydrophobicity or hydrophilicity). These three advantages constitute our central goal for using IMD as an educational tool, and regard the actions of protein unfolding as a practical educational tool to enhance and stimulate students’ undersatadning, and contextual skillsets through IMD.

#### 3) Folding Transition States of IMD

For this study, two IMD results were generated from the simulations of unfolding a protein: partially (complex) or totally unfolded (simple). The first group is a slightly or partially unfolded protein, where limited yet obvious distortions on the protein’s structure can be observed (complex structure: Fig.1a). The second group is an almost fully unfolded protein, where the protein has lost most of its secondary structure and becomes a coil from the amino acid sequence (complex structure: Fig. 1b). Thus, as a starting model, it is best to examine how our interactive model can detect and distinguish these clearly different groups of outcomes for a problem that has multiple solutions, where labeling a unique outcome can be confusing. It is worth mentioning that traditional molecular dynamics simulations do not generate these solutions. Instead, we explicitly invite a student with a molecular biology or biotechnology background to interact with the protein through IMD as much as possible before a threshold number of simulation steps is reached (here, we chose 100,000 steps). The only instruction the student has received is to nudge or stretch the protein and lysozyme here, starting from any set of residues that they prefer.

**Fig. 1.**
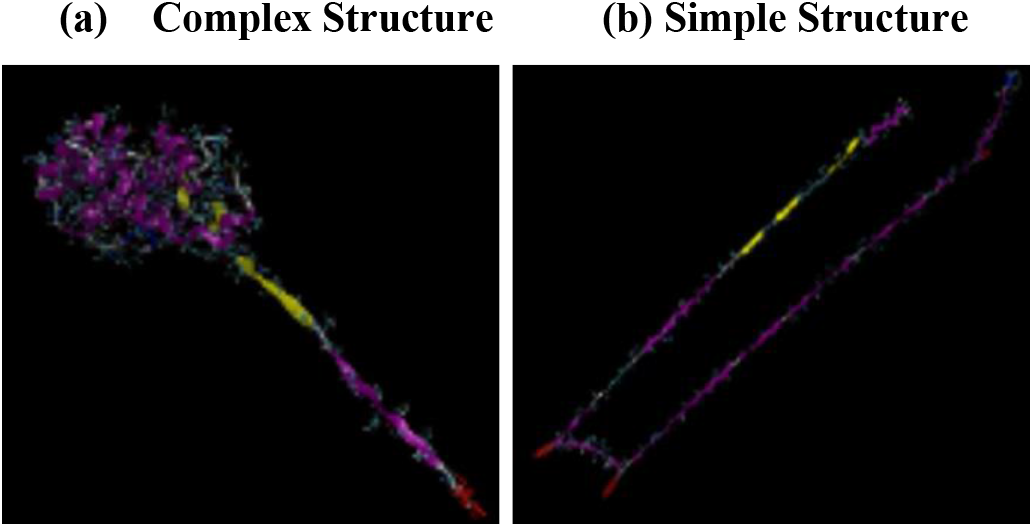
Folding transition states of two IMDs: (a) Largely Unfolded, and (b) Unfolded-Torn.

Therefore, data from the two simulations in Fig. 1 exemplifies a smaller set of outcomes we expect to see in a standard virtual learning environment (e.g., lab or classroom). Only the energy outputs were analyzed for this proof-of-concept study, which did not consider the simulation’s resulting molecular trajectories. The rationale for this is based on our objective to provide feedback to students on the fly, including students who may only have access to limited computing resources. Thus, making accurate enough predictions with a smaller data set is more desirable than a full-scale analysis of all molecular data in the background, which can be computationally intensive.

Among the energy outputs of each simulation, we choose to analyze the system’s kinetic energy (KE), potential energy (PE), and action (*S*), which is equivalent to the subtraction of PE from KE [31]. KE and PE are good features of our HAT context because KE reflects the overall motion of the system’s atoms in terms of the speed of individual atoms. Thus, changes in KE allow our model to detect HCI *if a student applies artificial forces to unfold the protein*. Alternatively, PE measures the interactions within the protein, and changes in PE inform us of intrinsic protein conformations in response to external forces. This can be considered after the student’s force (KE) *AI’s adaptive response to this force* (PE). Lastly, the action (S) is a quantity to be minimized in classical mechanics, which drives most of the motions observed in IMD. The minimization of *S* between two physical states generates an equation of motion for the system to obey. Changes in *S* offer insights into whether *a state transition of the protein in response to the student’s interaction is driven solely by nonequilibrium conditions created by the students or whether it relaxes responding to the external force for reaching certain new intermediates*. Thus, KE, PE, and *S* appear as appropriate starting features for modeling IMD within the context of HAT-HCI.

## II. Nonlinear Dynamical Systems Methods (NDS)

To better understand the dynamical characteristics of two distinct IMDs, we applied two nonlinear dynamical systems (NDS) methods in order: (1) attractor reconstruction and (2) recurrence quantification analysis (RQA). Using attractor reconstruction, we explore the changes in ‘stability’ (the Largest Lyapunov Exponent – LLE [32]–[34]) while using RQA, we examine ‘predictability’ (percent Determinism – DET [7], [35]) of IMD trajectories as a signature for HAT. Each IMD as a dynamical system exists in a state space (multidimensional space), which is an abstract construct depicting the range of behavior open to the system and is constrained by the degrees of freedom available to the elemental components of the system [36]. A system’s location in the state space is often determined through self-organization, which occurs when a system is attracted to a preferred state out of many potential states (i.e., the system prefers certain conditions and behaviors). These preferred states are called “attractors,” determined by the interactions between the system’s components and its sensitivity to external conditions [37]. The presence of those attractors is related to both the predictability of the system’s behavior and its response to novel situations. More robust attractors have more stable but less deterministic/predictable behavior. In this study, the stability of IMD is the maintenance of the molecular interaction patterns over time. These interactions can change over time based on the degree of variability (a transition process) in the interaction dynamics. Variability is needed for changes in interaction and to achieve the right balance of stability and flexibility of movement [38]. Thus, the more robust attractors’ behavior has less variability, and their adaptation can be problematic during unexpected changes in the task environment.

### A. Attractor Reconstruction: The largest Lyapunov (λ)

λ (“lambda”) is estimated by a process called “attractor reconstruction,” wherein scalar time series are unfolded into the requisite number of dimensions in order to represent the dynamical system (attractor) that generated the time series [39], [40]. To understand and characterize the magnitude of dynamical system stabilization on the attractor, the divergence of two nearby trajectories’ exponential rate can be defined based on the values of λ [32], [33]: (-)λ (stability) means that the system stabilizes quickly and will rigidly stay in that equilibrium state; (+)λ (instability) means that the system destabilizes quickly and will erratically shift from state to state; λ values near zero (i.e., called parallel trajectories or metastability) are generally nimble and, though they might meander, will tend to perform well (as with λ of large magnitude, small negative λ tends toward stability, small positive λ tends toward instability [41]. In this study, attractor reconstruction was performed separately for each IMD to examine possible differences in the context of HAT-HCI.

### B. Recurrence Quantification Analysis: Determinism

One method designed to understand how complex IMD dynamics work is recurrence quantification analysis (RQA: [7]) developed under the NDS: [42]. It has been applied in various domains, including linguistic (categorical:[43], [44]) or motion data, continuous [45], [46]. We applied continuous RQA to KE, PE, and S from mouse-tracking simulated data to investigate the system’s predictability of real-time molecular interaction.

#### 1) Recurrence Plots

The basis of RQA is the Recurrence Plot (RP: [47]), which is an ideal tool for visualizing the temporal evolution of a dynamical system when a system revisits similar states by identifying all pairs of time points in which the system returns to the same state [48]. RPs visually represent how often a system revisits certain states or sequences over time. On RPs, the diagonal lines are the states of the system’s epochs of similar time evaluation [49]. From this definition, unpredictable (chaotic) processes have no (or very short) diagonals, while predictable (deterministic) processes have longer diagonals and fewer single, isolated recurrence points [35]. Univariate categorical and continuous recurrence analyses have successfully been used to detect changing protein movements across cells and viruses [50]–[52].

#### 2) Percent Determinism (DET)

Several metrics can be extracted from RPs of each IMD signal, including recurrence rate, DET (or predictability), the longest diagonal line, entropy, laminarity, trapping time, and longest vertical line. Because of the local focus of the study, we only considered a commonly used metric in the HAT context [53], [54], called percent DET, to characterize IMDs, calculated based upon the ratio of recurrence points forming diagonal lines to all recurrent points in the upper triangle of the RPs, see Formula 1 [55]:

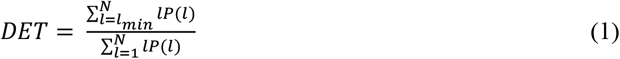

where “*l*” is the diagonal line length considered when its value is ≥ *l*_*min*,_ and *P*(*l*) is the probability distribution of line lengths. A *DET* of 0% means the time series never repeats, or the IMD is *highly unpredictable*, whereas a rate of 100% means the time series repeats perfectly, i.e., IMD is *highly predictable*. A highly deterministic (predictable) signal would be represented by several diagonal lines indicating that the signal repeats the sequences of IMD many times. Conversely, mildly deterministic IMD patterns rarely repeat a signal sequence and are represented by only a few diagonal lines [44]. In the IMD context, we interpret a percent DET value as *the percentage of IMD’s predictability by the interaction of a HAT*.

## III. Results

### A. Stability

To evaluate possible differences in each IMD stability, attractor reconstruction was performed separately for each of the IMD’s KE, PE, and S (N_full_= 100,000) and extracted their λ value. The exponential rate of divergence of two nearby trajectories on the attractor is measured by λ [32], [33] and characterizes the interaction stability of an IMD as either stable (λ < 0) or unstable (λ > 0) or metastable/ parallel trajectories (λ ≈ 0) [56]. In this study, attractor reconstruction was performed separately for largely unfolded and unfolded torn IMDs to examine possible differences in HAT-HCI dynamics. Reconstructed attractors for these two IMDs and their λs are illustrated in Fig. 2.

**Fig. 2.**
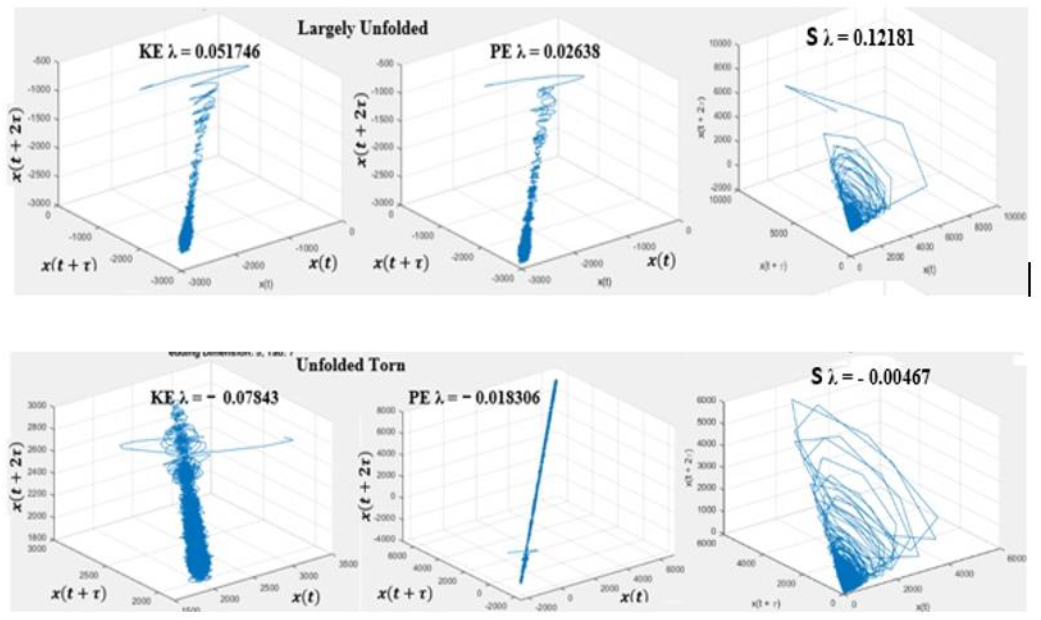
Reconstructed attractors from Largely Unfolded (top) and Unfolded Torn (bottom) IMDs’ KE, PE, and ACT functionals: a three-dimensional phase space as coordinates for the three-dimensional space [к(x), к(x+τ), к(x+2τ)]. The time delays are obtained via the average mutual information curve.

According to the visual representations of differently formed IMDs of Largely-Unfolded and Unfolded-Torn may differ in their temporal dynamics, λ estimates provide converging evidence of that observation. In Fig. 2 (top), while Largely Unfolded IMD’s λ_KE, PE, and S_ positive values (+) demonstrated more variability and less stability (unstable HAT-HCI behavior), Unfolded-Torn shows a stable IMD across all λ_KE, PE, and S_ negative values (–), which indicate too rigid HAT-HCI dynamics, Fig. 2 (bottom).

However, there is a clear distinction across KE, PE, and *S*’s temporal evolution within both IMDs. For instance, during a partially yet severely unfolded protein (Largely Unfolded), where distortions on the protein’s structure, reflecting on the loss of alpha helices and beta sheets, are substantial (see the visualization of previous figure - Fig. 1a), there was a decrease of λ values from KE to PE; an increment from PE to S. Changes in *S* from PE indicates that a state transition is spontaneous or driven by nonequilibrium conditions, such as student’s interference based on discovery curiosity, less stability but more adaptive system’s behavior.

On the other hand, unfolded torn IMD represents an almost-fully unfolded protein, where the protein has lost most of its secondary structure and becomes an almost 1-dimensional random coil that does not show a HAT-HCI adaptive dynamic behaviors increment from KE to PE, leading the system λ_S_ to metastable behavior. This can be interpreted as in unfolded torn IMD, the student’s too rigid (extremely stable) behavior led the computer to react with rigid interaction dynamics, which resulted in metastable system behavior (S) with negative λ, representing a less adaptive IMD paradigm.

### B. Predictability

Similarly, we applied RQA on the same IMDs’ KE, PE, and S measures to be consistent with attractor reconstruction. Illustrations of their RPs with their DET values are presented in Fig. 3. In both IMDs’ DET values, first, there is an increment from KE to PE and then a decrease from PE to S. Unfolded Torn has higher DET values (Fig. 3b) than Largely Unfolded DETs (Fig. 3a). However, the proportion of DET changes from KE-PE and KE-S for Largely-Unfolded IMD is higher than the Unfolded Torn. As an interpretation for this finding, Largely-Unfolded IMD can be a good simulation paradigm for system adaptation of IMD in which DET increment peaked from KE to PE - *the AI-tutor teammate’s interaction response to the student who demonstrated highly indeterministic interaction dynamics during the task*. However, we do not see this stochasticity in unfolded torn IMD. It means the system-level predictability was almost the same as the student’s unpredictable behavior. Whereas, in largely unfolded IMD’s system’s (S) DET shows that proportional change of DET from KE to S peaked regarding the computer’s predictable response (PE-DET) to the student’s highly unpredictable behavior (KE-DET). This can indicate AI’s adaptive behavior to the learner’s unpredictable behavior changes.

**Fig. 3.**
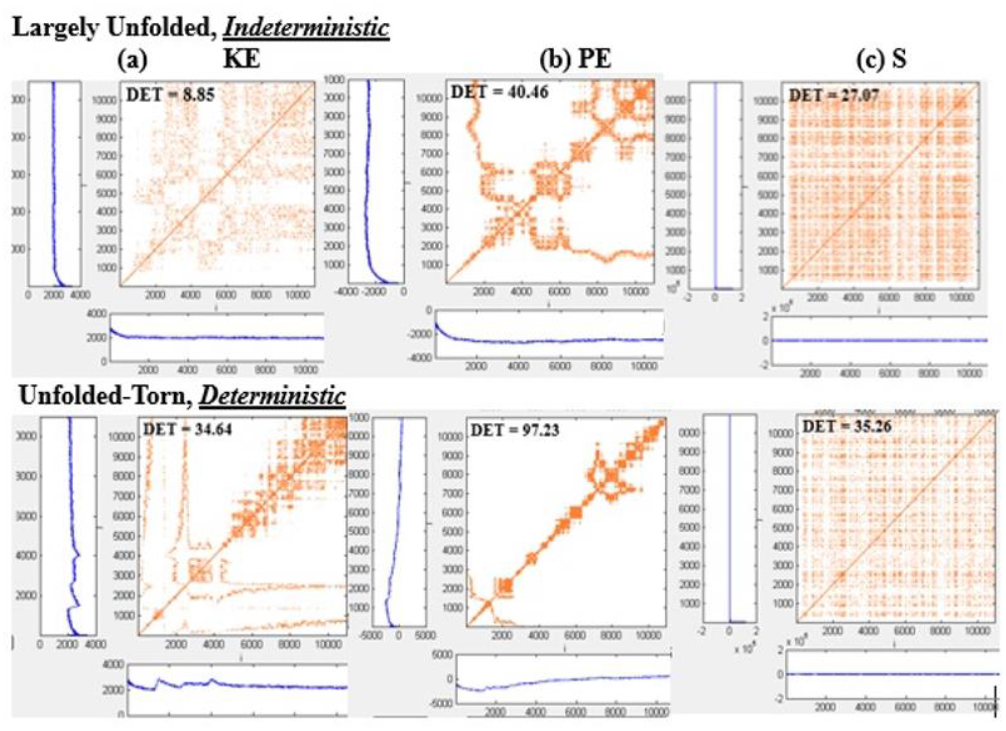
KE, PE, and S - RPs with their DET values of Largely Unfolded (a - Top) and Unfolded Torn (b - bottom) IMDs.

## IV. Discussion and Conclusion

Human-AI Teams (HATs) have become a vital instantiation of mixed teams in which humans and AI technology with a high level of role interdependency and heterogeneity work together to complete a common goal. Integrating AI as a teammate tutor into a virtual classroom environment can drive adaptive instruction and real-time constructive feedback to students, offering a possible conduit for fostering discovery curiosity in learners. However, integrating an AI as a teammate requires identifying signatures that track concomitantly with effective student(s) coordination and the dynamic task environment in which the system is subject.

Thus, for harnessing ‘discovery curiosity’ and learning, team coordination dynamics must be considered in developing the AI tutor and HAT system in the context of HCI. Because interaction recurrences between the system components depend on one another, the AI-tutor teammate must coordinate effectively with each student’s unique interaction behavior by maintaining the stability and predictability of the system to accomplish interactive learning tasks.

### A. NDS Modelling Outcomes and Discovery Curiosity

Each IMD’s stability (λ) and predictability (DET) derived from HAT-HCI endeavors distinguish between generic deprivation-driving and unique curiosity-driving interactions. The visualization and characterization of IMDs via attractor reconstruction and RPs and extracting λ and DET metrics, respectively, from them for probing variables indicate that the temporal evolution of IMDs can be determinants of AI-tutor teammate development.

One of the outcomes is the stability of IMD, i.e., the maintenance of the molecular interaction patterns over time. Maintaining IMD stability is needed as HAT adapts to the changes during the learning task, from routine to novel situations or vice versa. One of the IMD cases, Largely Unfolded, demonstrated an unstable coordination behavior at both energy (individual) and action (system) levels over time. This unstable behavior is not surprising because of the protein complexity, a partially yet severely unfolded protein, where distortions on the protein’s structure, reflecting the loss of alpha helices and beta sheets, are substantial. However, it is possible that incrementing the sample size in these types of protein forms can lead to more stable systems-level behavior because of the adaptation to the complex learning task. This example shows the idea of process-oriented training, perturbation, or coordination training [57], [58], in which either an all-human team or HAT can match coordination variability during the team task demands (anticipation of team member(s)’ needs about by coordinating the right information at the right time and place). Consequently, these findings verify the recent findings about process-oriented approaches to perturbation/ coordination training that can be associated with more adaptive HATs [58].

On the other hand, Unfolded Torn IMD represents a coordination dynamic that is too rigid due to the nature of the protein form, i.e., almost fully unfolded, where the protein has lost most of its secondary structure and becomes a coil from the amino acid sequence. Unfolded torn IMD represents a less adaptive IMD paradigm in which the student’s rigid (extremely stable) behavior led the computer to react with rigid interaction dynamics, which resulted in metastable system behavior (S) but still tends to be negative λ, representing stable behavior. In comparison to Largely Unfolded IMD, unfolded torn is a more accessible learning task for a student to follow via interacting with an AI tutor teammate. The Unfolded Torn IMD learning task is more relatable as a group task (homogeneous task roles) than a team task (heterogeneous task roles), Largely Unfolded IMD.

These interactions can change during the interactive learning task based on a transition process in the team coordination dynamics. In the previous HAT studies, we know that variability is needed to adapt to unexpected dynamic task environments to achieve the proper equilibrium of stability and flexibility of team coordination dynamics [53], [54]. Therefore, the second outcome, IMD predictability, showed that the largely unfolded IMD requires more flexible coordination dynamics to adapt to the complex protein unfolding than the unfolded torn. Unfolded-Torn’s extreme homogeneity requires a minimum level of predictability rather than flexibility.

### B. From Individual to Team Level Discovery Curiosity

In the previous curiosity studies, epistemic curiosity was investigated mostly individual level. However, this study explores ‘discovery curiosity’ not only individual level by using measures of KE and PE as representations of student and AI, respectively, but also team level via the S measure. Notably, the consistently obtained positive (‘+λ’ Largely Unfolded) and negative (‘-λ’) values of Unfolded Torn from KE, PE, and S measures can be an essential indicator to understand how coordination emerges as a property from individual taskwork to teamwork level (global-level). As indicated above, in a team task, ‘discovery curiosity’ to obtain new knowledge of taskwork and teamwork can be achieved by who knows what information and how it can be coordinated among the teammates (human or AI). It means that not only humans but also AI should have, at some level, ‘discovery curiosity,’ which alters maximizing the AI’s learning about its own taskwork and teamwork by interacting with other team members and its task environment in which the team is subject.

In this manner, if discovery curiosity requires information exchange across the team members to accomplish a shared task, then all the team members should have some level of discovery curiosity (more than some of its parts). This requires more in-depth future research on the PE measure as a representation of AI. We believe this will help develop more effective human-centered development of AI tutor teammates in the context of discovery curiosity, obtaining taskwork and teamwork knowledge, and learning.

### C. Implications of Human-AI Teaming

Stability and predictability show that a dynamic process-oriented task containing heterogeneous protein forms requires increased flexibility and less stable HAT interaction dynamics than homogeneous protein forms, e.g., unfolded torn IMD. Based on these findings, we have learned that a required level of team coordination dynamics is crucial for adaptive learning in complex IMD tasks. Thus, in HATs used in IMD tasks, we must build mechanisms to obtain an optimum level of team coordination dynamics between AI tutor teammate and learner in which the protein forms change the level of uniformity (homogeneous vs. heterogeneous). For future directions, the AI tutor teammate needs to be designed and tested under different levels of uniformity of IMDs to see how each HAT’s coordination dynamics evolve within teamwork and adaptive learning. In the long term, the results from the follow-up empirical studies will indicate the crucial characteristics needed of an AI-tutor teammate to interact on a peer-to-peer level with student teammates, which can advance the emergence of an AI tutoring system to support discovery curiosity, knowledge development, and applied learning in biotechnology.

## Acknowledgment

This material is based upon work supported by the National Defense Education Program (NDEP) for Science, Technology, Engineering, and Mathematics (STEM) Education, Outreach, and Workforce Initiative Programs under Grant No. HQ0034-21-S-F001. The views expressed in written materials or publications and/or made by speakers, moderators, and presenters do not necessarily reflect the official policies of the Department of Defense, nor does mention of trade names, commercial practices, or organizations imply endorsement by the US Government.

